# On-site integrated production of cellulases and β-glucosidases by Trichoderma reesei Rut C30 using steam-pretreated sugarcane bagasse

**DOI:** 10.1101/461012

**Authors:** Marcella Fernandes de Souza, Elba Pinto da Silva Bon, Ayla Sant’ Ana da Silvab

## Abstract

The high cost of commercial cellulases still hampers the economic competitiveness of the production of fuels and chemicals from lignocellulosic biomasses. This cost may be decreased by the on-site production of cellulases with the integrated use of the lignocellulosic biomass as carbon source. This integrated approach was evaluated in the present study whereby steam-pretreated sugarcane bagasse (SPSB) was used as carbon source for the production of cellulases by *Trichoderma reesei* Rut C30 and the produced enzymes were subsequently used for SPSB hydrolysis. An enzyme preparation with a high cellulase activity, of 1.93 FPU/mL, was obtained, and a significant β-glucosidase activity was achieved in buffered media, indicating the importance of pH control during enzyme production. The hydrolysis of SPSB with the laboratory-made mixture resulted in a glucose yield of 80%, which was equivalent to those observed for control experiments using commercial enzymes. Even though the supplementation of this mixture with external β-glucosidase from *Aspergillus awamori* was found to increase the initial hydrolysis rates, it had no impact on the final hydrolysis yield. It was shown that SPSB is a promising carbon source for the production of cellulases and β-glucosidases by *T. reesei* Rut C30 and that the enzyme preparation obtained is effective for the hydrolysis of SPSB, supporting the on-site integrated approach to decrease the cost of the enzymatic hydrolysis of lignocellulosic biomass.

## 1. Introduction

In the last few years, several industrial facilities aiming at the production of cellulosic ethanol have begun operating in different localities [1]. In Brazil, there are two facilities aiming at producing ethanol from sugarcane bagasse and straw, highly available feedstocks that amount to about 180 million tons of dry biomass per year [2, 3]. Nevertheless, this industry still faces several challenges, with economic studies showing that enzyme costs significantly affect the minimum ethanol selling price, resulting in a negative impact in the technology’s competitiveness in the biofuels market [4-6].

Industrial production of cellulases for lignocellulosic biomass hydrolysis is predominantly done by the companies Genencor (now part of DuPont) and Novozymes [7]. Even though these companies have succeeded in lowering the costs associated with the production of cellulases, techno-economic analyses suggest that the on-site production of enzymes may further contribute to the necessary cost decrease [8]. If the enzymes are produced at the same industrial site where the enzymatic hydrolysis of biomass will take place, there would be no need to concentrate, stabilize and transport the enzymatic cocktail before its ultimate use. Indeed, this approach has been shown feasible as cellulases have been already produced on site at a demonstration plant for the production of ethanol for sugarcane bagasse in Thailand [9]. This is especially relevant to countries like Brazil, whose continental sizes are obstacles to the transportation of off-site centralized enzyme production.

*Trichoderma reesei* is the most widely studied fungus for cellulase production and the most commonly used species for industrial production of these enzymes [10, 11]. *Trichoderma reesei* Rut C30, probably the best-known mutant, was selected because of its catabolite derepression and its high protein titers [12], which are about three times higher than those of the wild-type QM6a, selected by Mandels and Weber in the late 1960’s [13, 14]. However, its secreted enzyme mixture has typically been reported as unbalanced regarding the cellulase:β-glucosidase ratio (FPU:BGU), which may limit the enzymatic hydrolysis of cellulose into glucose [15-17].

To overcome this drawback, *T. reesei* cellulases are frequently supplemented with β-glucosidase from *Aspergillus* [18, 19], although the heterologous expression of β-glucosidase genes in *T. reesei* has also been studied [20, 21]. Nevertheless, few studies regarding the influence of the growth media on β-glucosidase production by *T. reesei* and the minimum FPU:BGU ratio necessary for an efficient hydrolysis have been carried out.

Several carbon sources, either soluble and insoluble, have been used for the production of cellulases by *T. reesei* [22, 23]. The choice of carbon source is a very important process parameter, since it has a significant influence on the cost of enzyme production [4] and on the final cellulase activity [22, 24]. Pretreated lignocellulosic biomasses have been shown to have inducing effects on cellulase secretion, depending on the chosen pretreatment, even resulting in higher β-glucosidase production in *T. reesei* [25]. This is a quite interesting scenario, as the pretreated feedstock to be hydrolyzed would be readily available for the on-site cellulases production, reducing even more the production costs [26]. Pretreatments based on the use of steam are considered effective for sugarcane bagasse, resulting in high glucose yields [27, 28]. These pretreatments remove the hemicellulose, yielding a more exposed cellulose fraction to the action of cellulases, and are already being used in industrial cellulosic ethanol facilities [29].

Therefore, the aim of the present work was to obtain a balanced cellulase and β-glucosidase enzyme preparation from *Trichoderma reesei* Rut C30 in an integrated production approach, using steam-pretreated sugarcane bagasse (SPSB) as carbon source in comparison to lactose and wheat bran. The laboratory-made enzymes, together with commercial cocktails, were compared for their adequacy in hydrolyzing SPSB and the influence of different FPU:BGU ratios in the hydrolysis process was assessed.

## 2. Materials and methods

### 2.1. Sugarcane bagasse and pretreatment conditions

Sugarcane bagasse was provided by Centro de Tecnologia Canavieira (CTC), Piracicaba, São Paulo, Brazil and pretreated at the Department of Chemical Engineering, Lund University, Sweden. Prior to steam pretreatment, the bagasse was impregnated overnight with 3% CO_2_ at 5 °C [27]. The impregnated material was steam-pretreated using a unit equipped with a 10-L reactor vessel at 220 °C for 5 min. Steam-pretreated sugarcane bagasse (SPSB) was characterized according to standard procedures [30] and its composition was 59.1% glucan, 4.0% xylan and 31.2% lignin.

### 2.2. Microorganisms maintenance, propagation and enzyme production

*Trichoderma reesei* Rut C30 (ATCC 56765) stock cultures were maintained on agar plates containing potato-dextrose agar. After incubation at 30 °C for seven days, these were observed as greenish sporulating cultures. To prepare the submerged culture inoculum, conidia were suspended in 5 mL of sterile water, and 1 mL of this suspension (10^6^ spores) was transferred to a 500-mL Erlenmeyer flask containing 100 mL of sterile Mandels medium [14]. The inoculum culture was incubated at 30 °C and 200 rpm for four days. For enzyme production, 30 mL of the mycelium suspension from the inoculum cultures was used to initiate growth in a 1-L flask containing 300 mL of growth medium as follows (in g/L): 0.3 urea, 1.4 (NH_4_)2SO_4_, 2.0 KH_2_PO_4_, 0.3 CaCl_2_, 0.3 MgSO_4_·7H_2_O, 6.0 yeast extract and 30 of either lactose (LAC), wheat bran (WB) or SPSB as carbon sources, plus 0.6% (v/v) corn steep liquor. Media were supplemented with trace elements as follows (in mg/L): 5 FeSO_4_·7H_2_O, 20 CoCl_2_, 1.6 MnSO_4_ and 1.4 ZnSO_4_. Media were buffered with 100 mM sodium phosphate buffer pH 6.0. Unbuffered fermentations with lactose as the carbon source were also carried out. The production cultures were incubated in an orbital shaker at 30 °C and 200 rpm up to seven days. Samples (10 mL) were taken every 24 hours for the measurement of filter paper (FPase) and β-glucosidase activities and the determination of pH.

*Aspergillus awamori* (2B.361 U2/1, classified by the Commonwealth Mycological Institute as an *Aspergillus niger* complex) maintenance and spore suspension preparation were performed as described for *T. reesei* Rut C30. Inoculum preparation was completed by two days of cultivation at 30 °C and 200 rpm in growth media containing (in g/L): 1.2 NaNO_3_, 3.0 KH_2_PO_4_, 6.0 K_2_HPO_4_, 0.2 MgSO_4_·7H_2_O, 0.05 CaCL, 12 yeast extract and 30 native sugarcane bagasse as carbon source. For enzyme production, a total volume of 30 mL of mycelium suspension, obtained from inoculum cultures, was used to initiate growth in a 1-L flask containing 300 mL of fresh medium. The culture was incubated in an orbital shaker at 30 °C and 200 rpm up to seven days.

### 2.3 Enzyme assays and analysis methods

The activities of total cellulase (filter paper activity, FPase) and β-glucosidase were determined according to standard IUPAC procedures [31]. One unit of FPase (FPU) corresponded to the release of 1 micromole of glucose per minute using an enzyme dilution providing 2 mg of glucose after 60 min assay reaction. One unit of β-glucosidase (BGU) was defined as the amount of enzyme that released 1 μmol of glucose in 1 min at 50 °C. Glucose concentrations for the β-glucosidase assay were measured using a biochemical analyzer (YSI 2700 Select, Yellow Springs, Ohio, USA).

### 2.4 Enzymatic hydrolysis of SPSB

Hydrolysis experiments were performed in 250-mL Erlenmeyer flasks containing 100 mL of the reaction mixture in sodium citrate buffer (50 mM), pH 5.0 and incubated at 50 °C and 200 rpm in an orbital shaker. The reaction mixtures contained 2.5% (w/v) SPSB, which had previously been dried overnight at 60 °C, plus the relevant enzyme pool. Enzymes, either produced in this work or commercially prepared (GC 220 and Spezyme^®^ CP from Genencor^®^), were loaded into the reaction mixture with 15 FPU/g of dry biomass.

The supernatants from the 7^th^ day of cultivation were used as source of the enzymes produced on lactose, wheat bran or SPSB. For the experiments with different FPU:BGU ratios, a concentrated supernatant of the 7^th^ day of cultivation in unbuffered lactose medium was used for the enzyme preparation with the 1:0.06 FPU:BGU ratio. A concentrated supernatant of the 7^th^ day of cultivation in buffered lactose medium was used for the enzyme preparation with the 1:0.38 FPU:BGU ratio. This supernatant was supplemented with *A. awamori* enzymes, produced as described in item 2.2, for obtaining mixtures with FPU:BGU ratios of 1:1 and 1:3. Enzyme concentration was performed in an Amicon 8200 stirred cell filtration module (Millipore, Billaica, USA) with a 10kDa membrane.

Monosaccharides were analyzed using an HPLC system equipped with a refraction index detector (RI-2031 Plus, Jasco Co., Japan) using an Aminex HPX-87P column (7.8 mm I.D. × 30 cm, Bio-Rad, USA) with a Carbo-P micro-guard cartridge (Bio-Rad, USA). Deionized water was used as the mobile phase at a flow rate of 1 mL/min at an 80 °C column temperature. Glucose concentrations were also measured using the Biochemistry Analyzer YSI 2700 Select. Hydrolysis experiments were carried out in triplicate.

Hydrolysis yields were calculated according to the equation [1]:

1

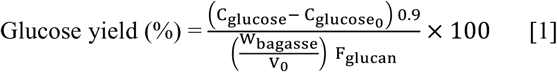

Where C_glucose_ is the glucose concentration in the hydrolysates (g/L); 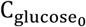 is the initial glucose concentration in the hydrolysis assay; W_bagasse_ is the total weight of bagasse in the hydrolysis assay (g); V_0_ is the initial volume of liquid (L); F_glucan_ is the initial mass fraction of glucan in SPSB.

## 2.5 Statistical analysis

The results from the triplicate experiments were compared using the Fisher’s LSD test (p ≤ 0.050) via Statistica^®^ (version 7.0). Error bars were plotted based on the standard deviations obtained from the experimental replicates.

## 3. Results and Discussion

### 3.1 Effect of different carbon sources and pH control on the production of FPase and β-glucosidase by *Trichoderma reesei* Rut C30

*Trichoderma reesei* was cultivated with steam-pretreated sugarcane bagasse (SPSB) as carbon source to assess the viability of using this material for enzyme production in an on-site integrated approach. Two other carbon sources, lactose and wheat bran, were used for comparison in simultaneous cultivations. Figure 1 shows FPase and β-glucosidase production obtained in buffered media containing the different carbon sources.

**Fig 1.**
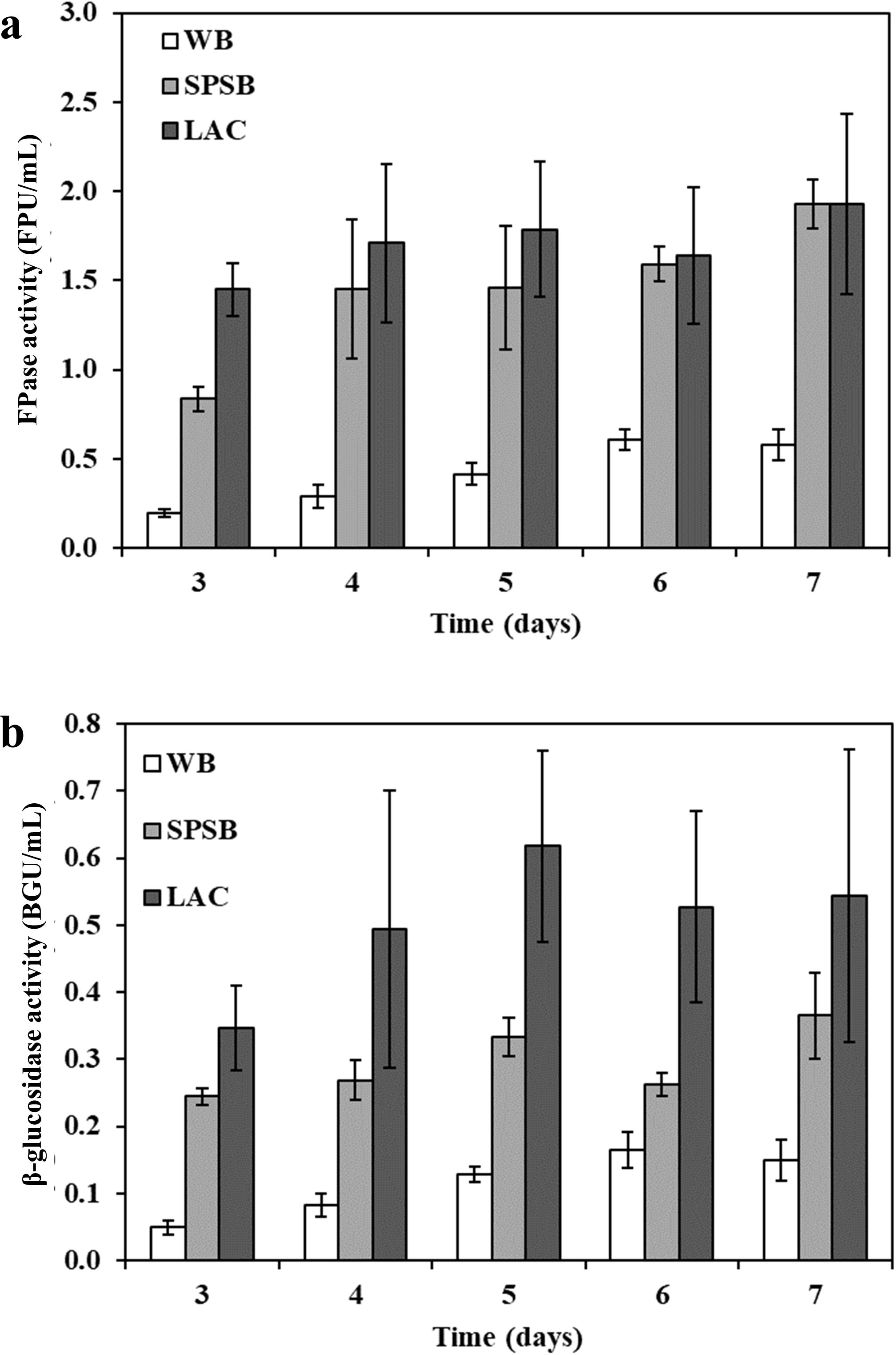
Time courses for FPase (a) and β-glucosidase (b) production by *T. reesei* Rut C30 in buffered media containing wheat bran (WB), steam-pretreated sugarcane bagasse (SPSB) or lactose (LAC) as carbon source

A maximum FPase activity of 1.93 FPU/mL was achieved when *T. reesei* was cultivated on either lactose or SPSB as carbon source and a 70% lower activity was measure for the cultivation on wheat bran. The low cellulose content of wheat bran, of around 11% (w/w) [32], in comparison to the SPSB’s cellulose content, of 60% (w/w), could be associated to the observed poor enzyme production. Even though wheat bran has been reported as beneficial for cellulase production by *T. reesei* in submerged fermentations, it is usually used as a complement to other carbon sources [33, 34].

Overall, lactose was the best carbon source for cellulase production, with peak activities reached between the 4^th^ and 5^th^ days of cultivation. This finding confirms the ability of lactose to induce cellulase production in *T. reesei* [35, 36]. Nevertheless, SPSB proved to be a promising carbon source for cellulase production with only a slightly lower productivity, reaching peak activities between the 6^th^ and 7^th^ days of cultivation. Maximum FPase activities reached in the cultivations with either lactose or SPSB showed no statistical difference (p<0.05), although the β-glucosidase activity achieved with SPSB was lower than that for the lactose medium (0.37 and 0.62 BGU/mL, respectively).

Several works have reported *T. reesei* as an ineffective microorganism for breaking down lignocellulosic substrates into glucose molecules due to its low β-glucosidase production [15-17]. However, it has also been reported that the amount of β-glucosidase in the culture filtrate of *T. reesei* Rut C30 may depend on the pH of the culture [37-40]. In this work, it was possible to obtain enzyme pools with rather significant β-glucosidase content, up to 0.62 IU/mL, in buffered medium containing lactose with an initial pH of 6.0 and a final pH of 5.8.

To better understand the effect of pH control in the production of β-glucosidases by *T. reesei* Rut C30, this fungus was cultivated in unbuffered media containing lactose. It was observed that the pH dropped to 2.5 and plateaued at this value from the 3^rd^ to the 7^th^ cultivation day. Under these conditions, the peak β-glucosidase activity corresponded to 0.02 BGU/mL, suggesting that low pH is an adverse condition for the enzyme production or stability. Interestingly, cellulase production was independent of pH control and a similar FPase activity, of 2.0 FPU/mL, was measured in both the lactose unbuffered and buffered medium.

Juhász et al. [38] attributed the higher β-glucosidase activity levels in buffered medium to the presence of maleic acid in the buffering system rather than to the actual culture pH. However, in the present study using phosphate buffer, with no maleic acid addition to the cultivation media, similar results were observed. Li et al. [39] and Ferreira et al. [40] also observed a higher β-glucosidase production by *Trichoderma reesei* when pH was controlled with succinate buffer or continuous adjustment with acid and base addition, corroborating that pH plays an important role in β-glucosidase production by *Trichoderma reesei*. A recent study evaluated the relationship between pH and gene regulation in *T. reesei* [41]. The authors found that the expression of genes encoding the major cellulases in *T. reesei* was not significantly affected under different pH conditions, which is in good accordance with the results reported in the present work. However, a gene encoding a candidate β-glucosidase displayed higher expression levels at pH 6 than at pH 3. Moreover, a mutant strain that behaved as if grown in acid conditions had a lower expression of two genes identified as encoding candidates for β-glucosidases when compared to the parental strain. Therefore, pH values higher than 3 seem to up-regulate the genes encoding β-glucosidase in *T. reesei*, resulting in a higher production of this enzyme with no effect on the production of FPase.

## 3.2 Effectiveness of enzymes produced with different substrates for SPSB hydrolysis

The hydrolytic efficiency of the culture supernatant of *T. reesei* grown in SPSB was evaluated in standard hydrolysis experiments alongside those obtained in the cultivations with lactose (LAC) or wheat bran (WB) and commercial cellulases. Table 1 presents the hydrolysis yields achieved with each enzyme preparation.

Previous reports indicate an advantage in using enzymes produced on the same substrate that would be further hydrolyzed [24, 42]. The present study, however, reported similar hydrolytic performance for enzymes produced on either lactose, wheat bran or SPSB (p>0.05). Our data are in good agreement with those of Jørgensen and Olsson [43], who reported that the hydrolysis yields for pretreated spruce were not affected regardless of the use of enzymes produced on different substrates, although the activity profile of the enzyme preparations could vary.

**Table 1.** Glucose yields (expressed as percentage of theoretical glucan content) after 6, 24 and 48 hours of hydrolysis of 2.5% SPSB using 15 FPU/g biomass. Enzymes were produced on different carbon sources, corresponding to: steam-pretreated sugarcane bagasse (SPSB), wheat bran (WB), LAC (lactose). Numbers shown in parenthesis indicate the FPU:BGU ratio of each enzyme preparation.

| Enzyme | Glucose yield (%) |  |  |
| --- | --- | --- | --- |
|  | 6 h | 24 h | 48 h |
| **SPSB (1:0.45)** | 23.4 ± 1.3 | 62.4 ± 3.2 | 79.6 ± 3.0 |
| **WB (1:0.50)** | 23.5 ± 0.1 | 63.2 ± 2.2 | 76.6 ± 3.0 |
| **LAC (1:0.76)** | 25.5 ± 0.1 | 61.0 ± 0.1 | 77.2 ± 5.6 |
| **GC 220 (1:0.78)** | 30.9 ± 0.9 | 68.9 ± 0.9 | 76.9 ± 0.6 |
| **Spezyme CP (1:0.97)** | 31.4 ± 1.3 | 71.4 ± 2.6 | 85.4 ± 3.4 |

When comparing the enzyme preparation produced on SPSB with the commercial preparations, it was observed a slightly lower efficiency of the SPSB enzyme mixture during the first 6h of hydrolysis (p<0.05); after 24 h, the SPSB mixture showed statistically similar performance to the GC 220 preparation, but the glucose yield obtained with it was still lower than that for Spezyme CP. Nevertheless, the 48h glucose yields for the SPSB preparation and that for the commercial enzymes (p>0.05), were similar, even though the SPSB preparation had the lowest FPU:BGU ratio.

Some studies using the integrated enzyme production approach with *T. reesei* did not report hydrolysis yields [22], not allowing a proper comparison of the results, as similar FPase activities may not directly correspond to similar hydrolysis abilities. Moreover, the reported results usually indicate low glucose yields for cellulase production from lignocellulosic biomasses [41, 46], unless when using a β-glucosidase supplementation [41]. Therefore, the results obtained in the present study, where SPSB was used for on-site enzyme production and no β-glucosidase supplementation was necessary, compares favorably to previous works.

In the hydrolysis experiments, the lowest FPU:BGU ratio evaluated corresponded to 1:0.45, resulting in a final glucose yield of almost 80%, which is considered satisfactory for cellulose-to-glucose conversion. This FPU:BGU ratio is much lower than the ratios usually reported as necessary for an efficient hydrolysis, of at least 1:1 [16, 18, 44]. In order to better understand the influence of the FPU:BGU ratio on the hydrolysis of SPSB, a *T. reesei* preparation with an FPU:BGU ratio of 1:0.38, produced on lactose, was supplemented with an *Aspergillus awamori* enzyme pool produced on sugarcane bagasse containing high levels of β-glucosidase (8.2 BGU/mL) and almost null FPase activity (0.06 FPU/mL), such that the final blends showed FPU:BGU ratios of 1:1 and 1:3. Additional hydrolysis assays using enzymes produced in an unbuffered lactose medium, with a FPU:BGU of 1:0.06, were simultaneously carried out. In these experiments, the SPSB concentration was increased to 7.5% (w/v) to enhance a possible detrimental effect of cellobiose accumulation in the final yields. The time course for glucose accumulation in experiments using the enzyme preparations with different FPU:BGU ratios are shown in Figure 2.

**Fig 2.**
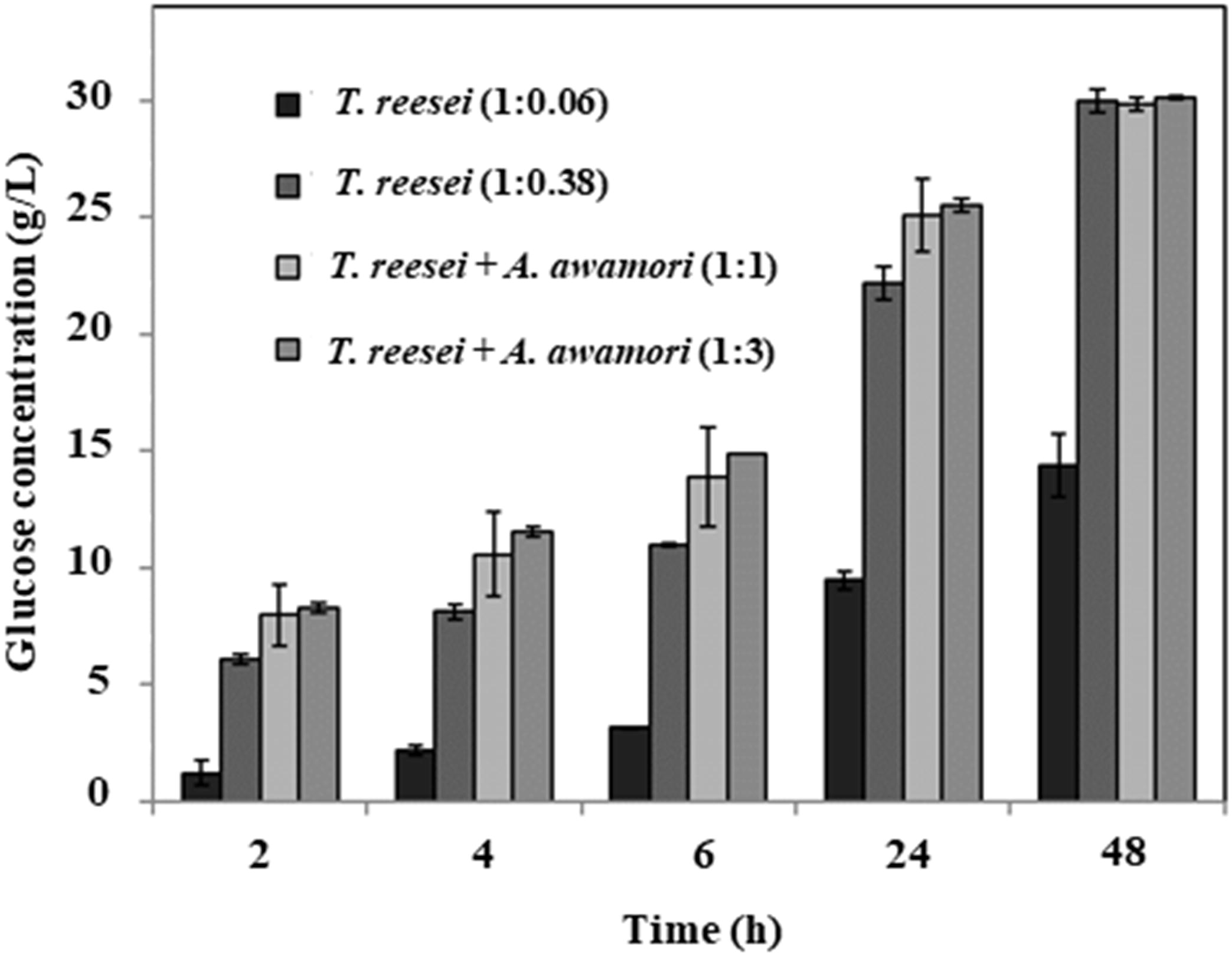
Glucose concentrations for the hydrolysis of 7.5% (w/v) steam-pretreated sugarcane bagasse (SPSB) using enzymes preparations secreted by *T. reesei* Rut C30 in unbuffered lactose medium (1:0.06), buffered lactose medium (1:0.38) and the latter supplemented with *Aspergillus awamori* enzymes (1:1 and 1:3). Numbers in parenthesis represent the FPU:BGU ratio

The use of an enzyme preparation with an FPU:BGU ratio of 1:0.38 resulted in a final glucose concentration two times higher than that obtained when using its unbuffered counterpart, suggesting that the use of a buffered medium results in the production of a more adequate cellulosic complex for the hydrolysis of SPSB. As expected, a cellobiose accumulation of 8.3 g/L was observed in the hydrolysis using the enzyme with an FPU:BGU ratio of 1:0.06, which most likely inhibited the action of cellulases.

Interestingly, the preparations with FPU:BGU ratios of 1:0.38 or higher gave similar glucose yields after 48 h of hydrolysis (p>0.05). Nevertheless, hydrolysis rates up to 24 h were higher in the experiments with an increased β-glucosidase activity (p<0.05), suggesting that a small supplementation could be beneficial, especially in experiments with higher solids loadings. This result is in good agreement with those obtained by Pallapolu and collaborators [45], who reported a beneficial effect of β-glucosidase supplementation only up to 24 h of hydrolysis for substrates poor on xylan, such as SPSB. For xylan-rich substrates, the authors found higher final hydrolysis yields with the supplementation and attributed it to accessory enzymes present in the β-glucosidase preparations, such as xylanases.

In the present study, an FPU:BGU ratio as low as 1:0.38 was enough for the hydrolysis of SPSB in low solids loading conditions (7.5% w/v). A similar result was obtained by Pryor and Nahar [46] with acid-treated corn cob and stover; the authors found that a 1:0.2 FPU:CBU (corresponding to a 1:0.4 FPU:BGU in the present study) was sufficient for the hydrolysis of these materials in low solids loadings. Therefore, we concluded that the necessity of β-glucosidase supplementation for biomass hydrolysis using the enzyme preparation produced by *T. reesei* Rut C30 may be lower than it is usually reported, and that SPSB was susceptible to hydrolysis using enzymes produced by *T. reesei* without supplementation. Future studies should evaluate the minimum β-glucosidase loading necessary for achieving high hydrolysis rates and yields in conditions of high solids loadings in order to assess the real necessity for this supplementation and guide the formulation of enzymes mixtures produced on-site. Indeed, the identification of the correct amount of cellulases and β-glucosidase would be valuable to the cost reduction of the integrated enzyme production and biomass hydrolysis processes. Even if a supplementation is found to be necessary, the present study indicates that the on-site integrated production of β-glucosidases would also be possible from *Aspergillus awamori* cultivation on sugarcane bagasse.

## 4. Conclusions

This study showed that *T. reesei* Rut C30 is able to produce a balanced enzyme preparation for the hydrolysis of SPSB when grown in a buffered medium. With this strategy, it was possible to exclude, or at least greatly reduce, the need for β-glucosidase supplementation, decreasing the relative cost of enzymes in the hydrolysis process. The effectiveness of SPSB as a carbon source to produce *T. reesei* cellulases and β-glucosidases is advantageous considering the on-site integrated enzyme production approach, as this substrate would be promptly available in the sugar mill, reducing the logistical costs for substrate and enzyme transportation.

## Acknowledgments

The authors would like to thank Dr. Clarissa Perrone for providing the samples of steam-pretreated sugarcane bagasse.

## Funding

This work was supported by the Brazilian National Council of Technological and Scientific Development (CNPq) and the Research and Projects Financing Agency [FINEP 1421-08].

